# Pexidartinib plus FLT3-directed CAR-Macrophage for the treatment of FLT3-ITD-mutated acute myeloid leukemia in preclinical model

**DOI:** 10.1101/2024.09.27.615313

**Authors:** Fan Chen, Min-Hong Lv, Chun-Wei Li, Yi-Qiong Zhang, Ze-Xiao Wei, Zhi Yuan, Jian-Yong Cao, Xinming Yang, Jing-Bo Xu, Li Wang, Bai-Liang He

**Author notes:** **Correspondence: Bai-Liang He, Ph.D.**, Address: Guangdong Provincial Engineering Research Center of Molecular Imaging, The Fifth Affiliated Hospital, Sun Yat-sen University, Zhuhai, Guangdong, China; Guangdong-Hong Kong-Macao University Joint Laboratory of Interventional Medicine, The Fifth Affiliated Hospital, Sun Yat-sen University, Zhuhai, Guangdong, China., **Jing-Bo Xu, M.D.**, Address: Department of Hematology, The Fifth Affiliated Hospital, Sun Yat-sen University, Zhuhai, Guangdong, China., **Li Wang, M.D. & Ph.D.**, Address: Department of Gynecology and Obstetrics, Perinatal Medical Center, The Fifth Affiliated Hospital, Sun Yat-sen University, Zhuhai, Guangdong, China. These authors contributed equally to this work (Fan Chen, Min-Hong Lv, Chun-Wei Li).

## Abstract

Acute myeloid leukemia (AML) is the most common type of acute leukemia in adults. Internal tandem duplication of FMS-like tyrosine kinase 3 (FLT3-ITD) mutations occur in about 25%-30% of AML cases and are associated with adverse prognosis. Recent advances indicate that M2-like leukemia-associated macrophages (M2-LAM) are highly infiltrated in the bone marrow of FLT3-ITD+ AML patients; however, the underlying mechanisms and therapeutic implications are still elusive. Herein, we reveal that conditioned medium from FLT3-ITD+ MOLM-13 AML cells polarized M2-LAM and impaired their phagocytic activities. Unexpectedly, co-culture of M2-LAM protected MOLM-13 cells from the treatment of FLT3 inhibitor quizartinib by activating their FLT3 signaling pathway. Pharmaceutically, FLT3/CSF1R dual inhibitor pexidartinb effectively suppressed M2-LAM, reduced leukemic burden, and prolonged the survival of MOLM-13-xenografted mice. To enhance the phagocytic activities of macrophages, FLT3-directed chimeric antigen receptor-engineered macrophages (FLT3L-CAR-Macrophage) were generated using FLT3 ligand (FLT3L) as the recognizing domain of CAR. Transfection of THP-1 monocytic cells-or umbilical cord blood mononuclear cells-derived macrophages with FLT3L-CAR-encoding mRNA enhanced their phagocytic activities to MOLM-13 cells in vitro. Consistently, FLT3L-CAR-Macrophage differentiated from FLT3L-CAR-expressing THP-1 cells effectively phagocytosed MOLM-13 cells in vitro, reduced leukemic burden and prolonged the survival of MOLM-13-xenografted mice. Importantly, treatment of pexidartinib resulted in upregulation and surface localization of FLT3-ITD protein in MOLM-13 cells, sensitized MOLM-13 cells to the treatment of FLT3L-CAR-Macrophage in vitro, and synergized with FLT3L-CAR-Macrophage to further reduce leukemic burden in MOLM-13-xenografted mice. Together, our data indicate that pexidartinib plus FLT3L-CAR-Macrophage could be a promising therapeutic strategy for the treatment of FLT3-ITD+ AML in preclinical model which warrants further investigation.

**Graphical Abstract:** *Pexidartinib plus FLT3L-CAR-Mφ for the treatment of FLT3-ITD+ AML in preclinical model. Pexidartinib suppressed AML growth, reduced M2-LAM, increased FLT3 surface expression and synergized with FLT3L-CAR-Mφ to target FLT3-ITD+ AML in vitro and in vivo*.

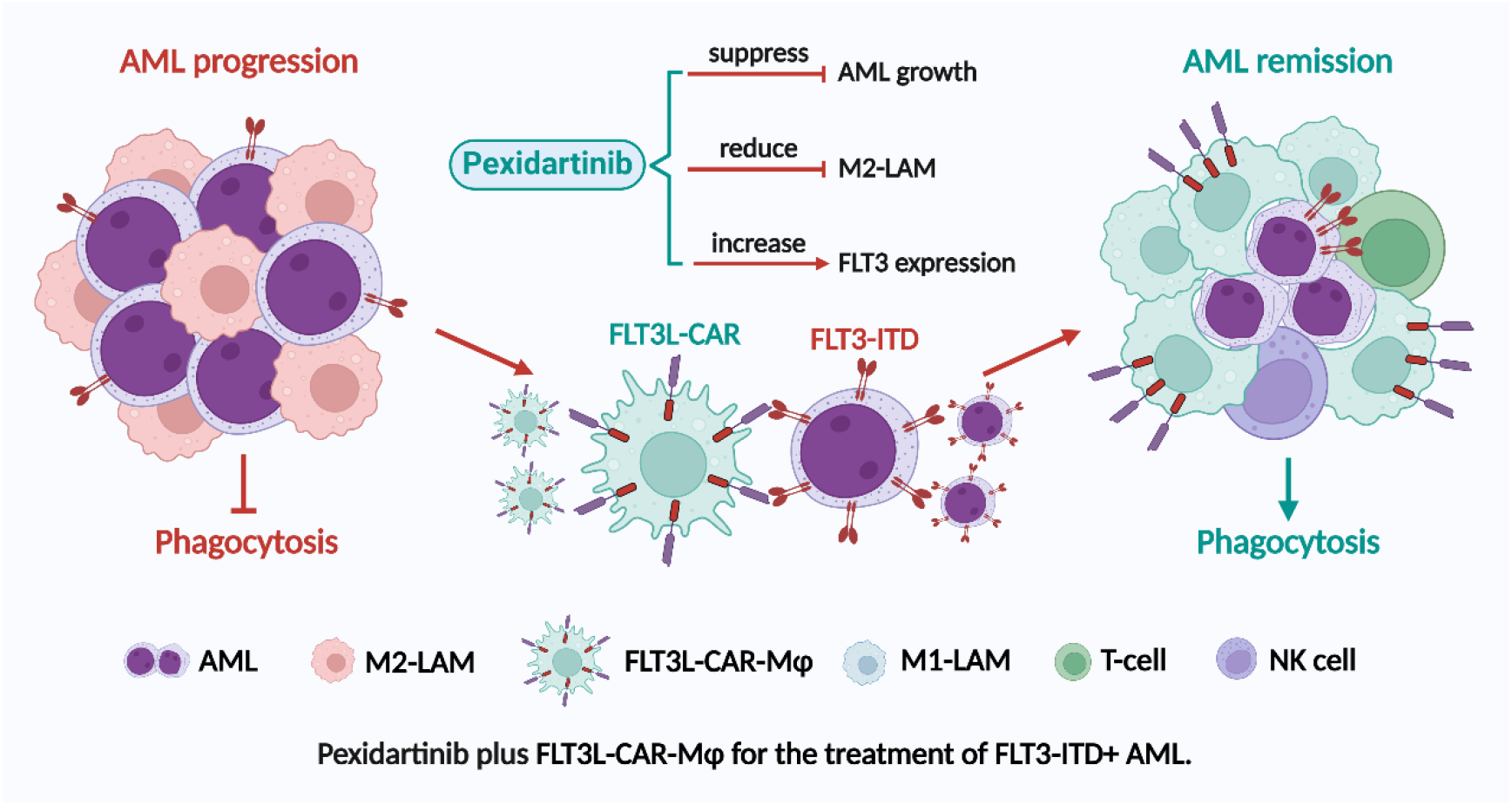

## Introduction

Acute myeloid leukemia (AML) is the most common type of acute leukemia in adults and characterized by abnormal expansion of myeloblasts in peripheral blood (PB) and bone marrow (BM).^1^ It is a heterogeneous disease with distinct cytogenetic and genetic features in individual patients.^2^ Internal tandem duplication of FMS-like tyrosine kinase 3 (FLT3-ITD) mutations occurs in about 25%-30% of AML cases and are associated with adverse prognosis.^3^ Though FLT3 tyrosine kinase inhibitors (FLT3-TKI) have been used in FLT3-ITD+ AML patients,^4-6^ the treatment outcomes are still not satisfactory.

We previously reported that the percentage of immunosuppressive M2-like leukemia-associated macrophages (M2-LAM) was increased in FLT3-ITD+ AML.^7^ Consistently, Brauneck and colleagues demonstrated that the frequencies of immunosuppressive TIGIT-positive M2-LAM were significantly increased in BM from FLT3-ITD+ AML and associated with adverse prognosis.^8^ However, the pathogenic roles and therapeutic implications of M2-LAM in FLT3-ITD+ AML are still elusive.

Here we show that FLT3-ITD+ AML cells polarized M2-LAM and impaired their phagocytic activities. M2-LAM protected FLT3-ITD+ AML cells from the treatment of quizartinib through activating FLT3 signaling. FLT3/CSF1R dual inhibitor pexidartinib reduced M2-LAM, upregulated FLT3, and synergized with FLT3-directed chimeric antigen receptor-bioengineered macrophages (FLT3L-CAR-Mφ) to target FLT3-ITD+ AML in vitro and in AML-xenografted mouse model.

## Materials and methods

### Cell lines

THP-1, MOLM-13 and MV4-11 cell lines are obtained from ATCC or DSMZ as previously described.^9^ Cells were cultured in regular medium (RM) consisting of RPMI medium supplemented with 10% FBS (Gibco, Thermo Fisher Scientific). Conditioned medium (CM) from MOLM-13 cells (CM-MOLM-13) was harvested and filtered (0.22 μm) as previously described.^10^ All cell lines were recently authenticated and confirmed as mycoplasma-free.

### Western blotting

MOLM-13 cells were collected and lysed (Cell lysis buffer, Beyotime) as our previously described.^7^ Equal amounts of protein (Pierce BCA Protein Assay Kit, Thermo Fisher Scientific) were separated by 10% SDS-PAGE gel, transferred to PVDF membranes, blocked with 5% milk in TBST, and then incubated with primary antibodies (Table. 1) overnight at 4°C. After washing in TBST three times, the membranes were incubated in HRP-conjugated secondary antibody (Beyotime, 1: 5000) at room temperature for 1 hour. Subsequently, the signals were detected by ECL (Immobilon Western HRP substrate, Merck Millipore) and visualized (iBright Imaging Systems, ThermoFisher). β-ACTIN was used as loading control. The relative protein expression was analyzed by ImageJ software after normalizing it to the corresponding ACTIN.

**Table 1.** Reagents and resources used in this study.

| **Name** | **Source** | **Identifier** |
| --- | --- | --- |
| **Western Blotting** |  |  |
| Anti FLT3 (8F2) Rabbit mAb (1:1000 dilution) | CST | 3462S |
| Anti phospho-FLT3 (Tyr591) Rabbit mAb (1:1000 dilution) | CST | 3461S |
| Anti STAT5 (D3N2B) Rabbit mAb (1:1000 dilution) | CST | 25656S |
| Anti phospho-STAT5 (Tyr694) Antibody (1:1000 dilution) | CST | 9351S |
| Rabbit anti human pan AKT (1:1000 dilution) | CST | 4685S |
| Anti phospho-Akt (Ser473) Antibody (1:1000 dilution) | CST | 9271S |
| Anti p44/42 MAPK (Erk1/2) Antibody (1:1000 dilution) | CST | 9102S |
| Anti phospho-p44/42 MAPK (Erk1/2) Rabbit mAb (1:1000 dilution) | CST | 4370S |
| Anti Stat3 (124H6) Mouse mAb (1:1000 dilution) | CST | 9139T |
| Anti phospho-Stat3 (Tyr705) (D3A7) Rabbit mAb (1:1000 dilution) | CST | 9145S |
| Anti-Beta-ACTIN (1:5000 dilution) | Sigma | A2228 |
| **Immunofluorescent staining** |  |  |
| Anti Myc-Tag (71D10) Rabbit mAb (1:200 dilution) | CST | 2278T |
| **Flow cytometry** |  |  |
| PE Mouse IgG1, $\kappa$ isotype Ctrl | Biolegend | 400111 |
| PE anti-human CD135 | Biolegend | 313305 |
| **Drug treatment** |  |  |
| Phorbol 12-myristate 13-acetate (PMA) | MCE | HY-1873 |
| Quizartinib | Beyotime | SC1049 |
| Sorafenib | APExBio | A3009 |
| Pexidartinib | TargetMol | T2115 |
| Midostaurin | TargetMol | T3211 |
| Crenolanib | TargetMol | T2677 |
| Gilteritinib | TargetMol | T4409 |
| **Software** |  |  |
| Flowjo | FLOWJO LLC | V7.6.1 |
| Prism | Graphpad | V8.0.1 |
| Image Lab | Bio-Rad | V5.2.1 |
| Photoshop | Adobe | V2017.0.0 |
| Living Image | Revvity | V4.4 |

### Real-time quantitative PCR

Real-time quantitative PCR (RT-qPCR) was performed as our previously described.^7,9^ Total RNA was extracted (NucleoZol, MACHEREY-NAGEL, Germany) from different cells and reversely transcribed (HiScript III RT SuperMix, Vazyme, China) into first strand cDNA. RT-qPCR was performed using SYBR Green Master Mix (Vazyme, China) by StepOnePlus Real-Time PCR System (QuantStudio 7 Flex, ABI). *GAPDH* was used as internal control (Table. 2). Relative gene expression was calculated using the 2^-^△△Ct method.

**Table 2.**
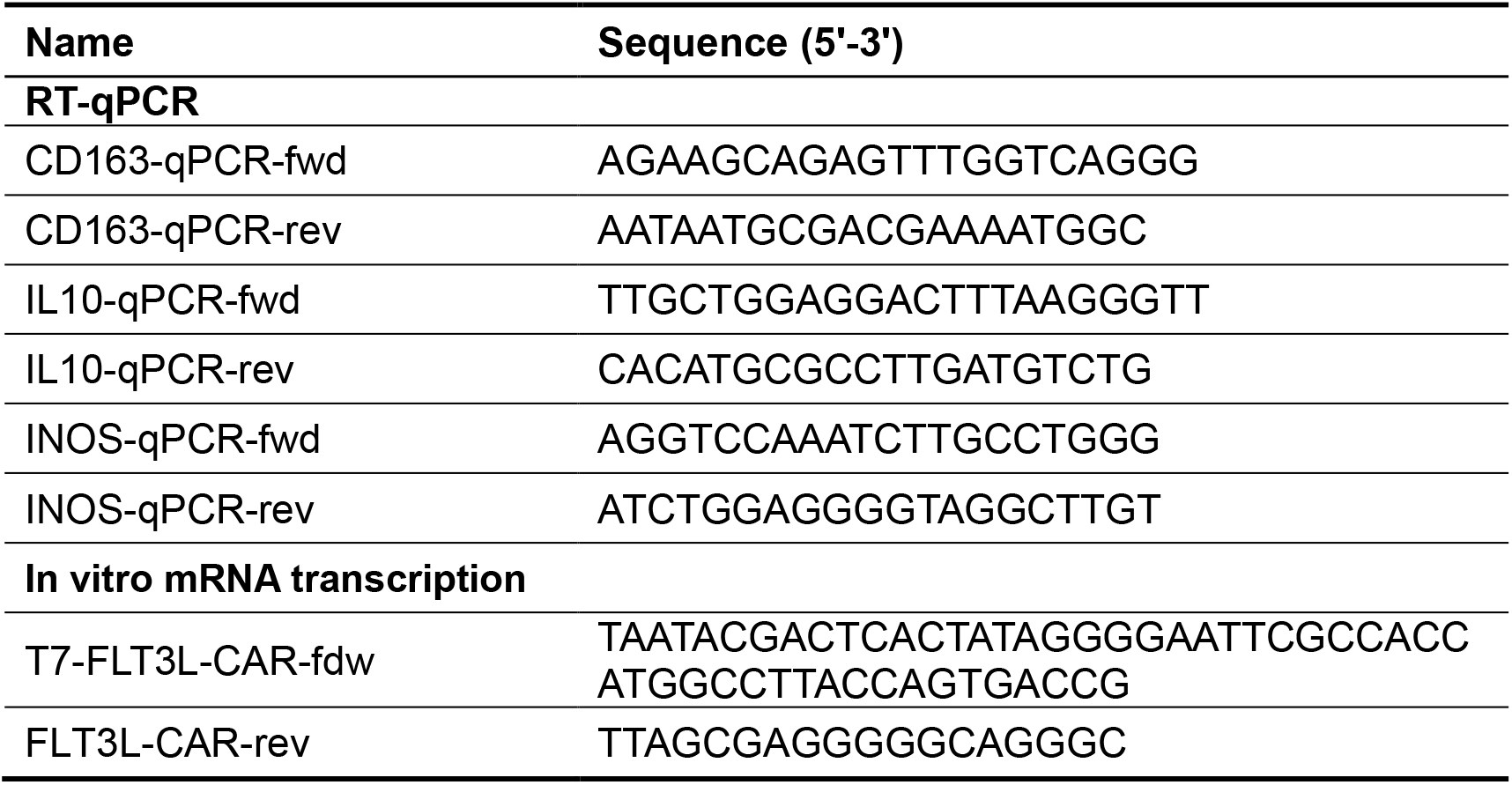
Primers and DNA sequence used in this study.

### Drug treatment

The anti-leukemic effects of candidate drugs were tested in leukemia cell lines as previously described. Briefly, leukemia cells (0.2 × 10^6^/mL, 100 μL) were plated in 96 well plates in triplicate and treated with defined concentrations of quizartinib (Beyotime) for three days in RM or CM from different macrophages. The effects of drug treatment were determined by cell viability assay (CCK8 Cell Viability Reagent, Beyotime). After normalization, the dose-response curves of quizartinib were generated by GraphPad Prism 8.0.1.

### Luciferase-based cell killing assay

THP-1 monocytic cells were seeded into 96-well plates (5,000 cells/well) and treated with PMA (20 ng/ml) to induce macrophage differentiation for 48 hours. MOLM-13 cells stably expressing luciferase (MOLM-13^luc^) were then added (5,000 cells/well) into the differentiated macrophages and treated with different concentrations of quizartinib for two days. 10 μl of luciferin (15 mg/ml, Beyotime, China) was added into each well and the luciferase intensity in each well was immediately measured using an Alpha Microplate reader (PerkinElmer). The viability of MOLM-13^luc^ cells was calculated after normalizing to the DMSO control and the dose-response curve of quizartinib was generated by GraphPad Prism 8.0.1.

### THP-1-derived macrophages

Immortal THP-1 monocytic cells were seeded in 6-well plate and treated with PMA (20 ng/ml) for 48 hours to induce macrophage differentiation as previously described.^11^ CM from macrophages (CM-Mφ) was harvested and filtered (0.22 μm) as previously described.^10^ Serum from primary AML bone marrow samples was collected as our previously described.^12^

### Generation of FLT3-directed CAR-Macrophages by mRNA transfection

The FLT3 ligand-based chimeric antigen receptor (FLT3L-CAR) were constructed (IGE Biotechnology, China) according to the previous reports with several modifications.^13,14^ Briefly, sequences of CD8α leader, FLT3 ligand (FLT3L), MYC tag, CD28 hinge, CD28 transmembrane, and intracellular CD3ζ were cloned into the pCDH-Puro-EV lentivirus vector (IGE Biotechnology, China) to generate pCDH-FLT3L-CAR. pCDH-FLT3-CAR vector was used as DNA template to amplify the full-length of FLT3L-CAR by PCR. T7 promoter was included in the forward primer (Table. 2) during the PCR assay. The PCR products were purified (PCR Clean Up Kit, Beyotime) and used as DNA template for in vitro mRNA transcription (HiScribe T7 ARCA mRNA Kit, NEB). FLT3L-CAR mRNA was capped with Anti-Reverse Cap Analog (ARCA) and tailed with poly-A to increase their translation efficiency and stability, respectively. In parallel, mRNA encoding for *EGFP* was generated as control. In vitro transcribed mRNA was purified by LiCl precipitation (NEB) and stored in -80°C until use. THP-1 monocytic cells were differentiated into macrophages in the presence of PMA for 2 days in vitro. Umbilical cord blood mononuclear cells (UCB-MNCs) were isolated by density gradient centrifugation (Ficoll-Paque, HyClone, Thermo Fisher Scientific) and differentiated into macrophages in the presence of GM-CSF (10 ng/mL) for 7 days in vitro. Differentiated macrophages were transfected with FLT3L-CAR mRNA (500 ng per well in 24-well plate) using Lipofectamine™ MessengerMAX™ reagent (Thermo Fisher Scientific).

### Generation of FLT3-directed CAR-Macrophages by lentivirus transduction

Lentivirus was packaged in 293T cells by co-transfecting lentiviral packaging plasmid pCMV-dR8.91, envelop plasmid pMD2.G, and pCDH-FLT3L-CAR transferring lentiviral vector as previously described.^9^ Immortal THP-1 monocytic cells were first transduced with mCherry-encoding lentivirus (Addgene #176016) and sorted by FACS (BD FACSaria FUSION). Subsequently, mCherry^+^ THP-1 cells were transduced with pCDH-FLT3L-CAR lentivirus by spinoculation in the presence of polybrene (8 μg/mL). Lentivirus transduced THP-1 cells were further enriched by puromycin (8 μg/mL) selection for three days in vitro. FLT3L-CAR-expressing THP-1 cells were expanded and treated with phorbol 12-myristate 13-acetate (PMA, 20ng/mL) for 2 days to differentiate into FLT3-directed CAR-Macrophages.

### Immunofluorescent staining

To detect the cellular expression of FLT3L-CAR (MYC-tagged), anti-MYC immunofluorescent staining was performed using primary macrophages- and THP-1-derived FLT3L-CAR-Mφ. Briefly, macrophages were seeded in glasses, washed with cold PBS, and fixed by 0.4% paraformaldehyde (PFA) for 15mins at room temperature. After blocking (5% BSA, 0.3% Triton X-100 in PBS) for 1 hour at room temperature, macrophages were stained with anti-MYC primary antibody (Table. 1) at 4°C overnight. After washing and incubating in FITC-conjugated secondary antibody, macrophages were counter stained with DAPI (Biosharp) and subjected to confocal imaging (Laser confocal microscopy, Zeiss 880).

### Microscopy-based phagocytosis assay

FLT3-ITD+ MOLM-13 cells labelled with CFSE (Biolegend) were used as target cells for phagocytosis assays. Primary macrophages-derived FLT3L-CAR-Mφ labelled with CM-Dil (1 μmol/mL, Yeasen) or mCherry+ THP-1-derived FLT3L-CAR-Mφ were used as effecter cells. MOLM-13 cells were co-cultured with FLT3L-CAR-Mφ (E: T=1: 4) for 12 hours. Then the supernatant was discarded and the attached macrophages were gently washed with PBS and subjected to fluorescent imaging (IX73 microscopy, Olympus) immediately. The phagocytic activity of FLT3L-CAR-Mφ was quantified by phagocytosis events per field (six fields per sample). Untransfected primary macrophages (CTL) or empty vector-transduced THP-1-derived macrophages (EV-Mφ) were used as control, respectively.

### Flow cytometry-based phagocytosis assay

After co-culture experiment and fluorescent imaging as shown above, the attached macrophages were harvested (0.25% Trypsin-EDTA, ThermoFisher) and subjected to flow cytometry (LSR Fortessa Cell Analyzer, BD Biosciences) analysis immediately.

The phagocytic activity of FLT3L-CAR-Mφ was defined by the percentage of MOLM-13-postive populations (CFSE+). All experiments were performed in triplicate.

### AML xenograft mouse model

NOD/ShiltJGpt-*Prkd*^cem26Cd52^*Il2rg*^em26Cd22^/Gpt (NCG) immunocompromised mice (defective in T, B, and NK cells) were purchased from Gempharmatech (China) and maintained in the standard specific-pathogen-free (SPF) condition in laboratory animal unit from the fifth affiliated hospital of Sun Yat-sen university. One day before xenotransplantation, 6-to 8-week-old NCG mice were conditioned by the treatment of busulfan (30 mg/kg). MOLM-13-luc cells (1 × 10^4^) stably expressing luciferase reporter (Addgene #46793) were injected through tail vein. In the next day, FLT3L-CAR-Mφ (5 × 10^6^) were injected through tail vein into five mice, randomly. Engraftment of MOLM-13 cells were confirmed by bioluminescence imaging (BLI) at one-week post injection. Leukemic burdens were examined by bioluminescence imaging at two-weeks post transplantation. Survival of animals from different groups was recorded.

### Statistical analysis

All experiments were performed in triplicates and data were presented as the mean ± standard error of the mean (SEM) unless otherwise specified. Numerical data were compared using the student’s t-test. Survival data was analyzed using the Kaplan-Meier method (log-rank test). All statistical analyses were performed using GraphPad Prism 8.0.1. *p* < 0.05 (*), *p* < 0.01 (**), and *p* < 0.001 (***) were considered statistically significant.

## Results

### Conditioned medium from FLT3-ITD+ AML cells polarized M2-LAM

It has been reported that the frequencies of immunosuppressive M2-like leukemia-associated macrophages (M2-LAM) are increased in the bone marrow from FLT3-ITD+ AML patients;^8^ however, the underlying mechanisms are still elusive. Herein, we first sought to study the interplay between FLT3-ITD+ MOLM-13 AML cells and macrophages in vitro. Conditioned medium (CM) from MOLM-13 cells polarized macrophages into M2-like phenotype as shown by increase of M2 markers CD163 and IL10 (**Fig. 1A-C**). The phagocytic activities of MOLM-13-educated M2-LAM were impaired compared to the control macrophages as shown by the reduction of double positive population (**Fig. 1D-F**). CM from M2-LAM (CM-M2-LAM) or co-culture with M2-LAM protected MOLM-13 cells from the treatment of FLT3 tyrosine kinase inhibitor (FLT3-TKI) quizartinib in vitro (**Fig. 1G-J**). Mechanistically, CM-M2-LAM induced increased phosphorylation of FLT3 and its downstream signaling including STAT5, STAT3, and AKT in MOLM-13 cells, comparing to the CM from control macrophages (**Fig. 1K-M**).

**Figure 1.**
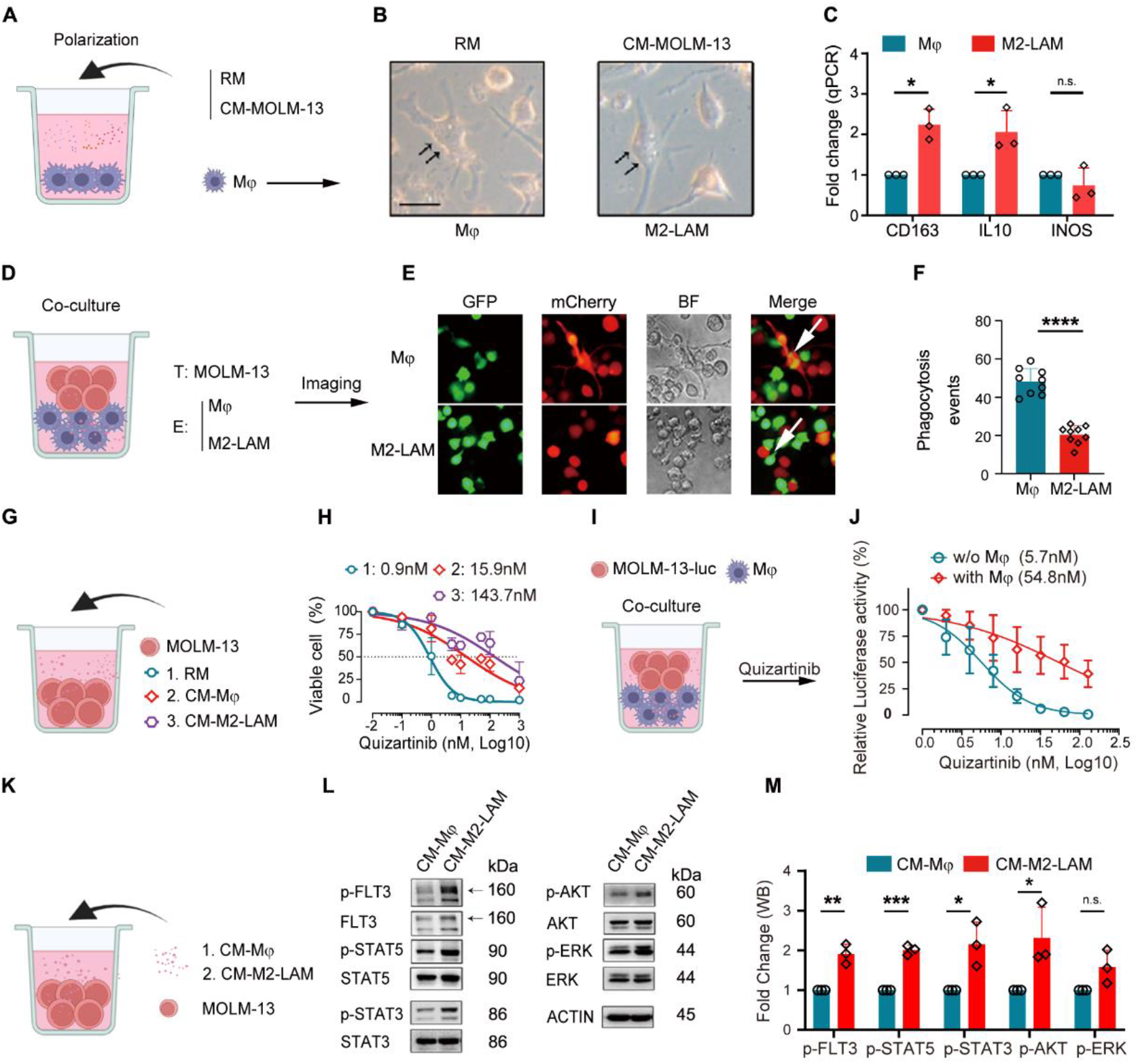
M2-LAM protected FLT3-ITD+ AML cells from the treatment of quizartinib. **(A-C)** Effects of regular medium (RM) and conditioned medium from MOLM-13 (CM-MOLM-13) cells on the polarization of THP-1-derived macrophages as shown by RT-qPCR of M2 (CD163 and IL10) and M1 (INOS) markers (C). **(D-F)** Detecting the phagocytic activity of control macrophages (Mφ) and MOLM-13-educated macrophages (M2-LAM) by fluorescent imaging (D, E). Phagocytic events per field was recorded and compared (F). **(G-H)** Dose response curve of quizartinib in MOLM-13 cells cultured with RM, CM from control Mφ (CM-Mφ), or CM from M2-LAM (CM-M2-LAM). **(I-J)** Dose response curve of quizartinib in MOLM-13-luc cells cocultured with THP-1-derived macrophages. The viability of MOLM-13-luc cells calculated after normalizing the luciferase activity in each well to the DMSO control. **(K-M)** Western Blotting showing the expression and phosphorylation of FLT3 and its downstream signaling pathway in MOLM-13 cells cultured in CM-Mφ and CM-M2-LAM.

### Targeting FLT3-ITD+ AML and M2-LAM by FLT3/CSF1R dual inhibitor pexidartinib

The above observations prompted us to test whether targeting M2-LAM may show therapeutic efficacy in FLT3-ITD+ AML. CSF1 receptor (CSF1R) is crucial for the differentiation of pro-tumoral and immunosuppressive M2-like tumor-associated macrophages (M2-TAM).^15^ As a novel FLT3/CSF1R dual inhibitor, pexidartinib (also known as PLX3397) was approved by FDA as monotherapy for symptomatic tenosynovial giant cell tumor (TGCT)^16^ and reproposed to target M2-TAM in a phase I clinical trial for soft tissue sarcomas (STS).^17^ Consistently, bone marrow serum from FLT3-ITD+ AML patients induced an increased proportion of M2-LAM which could be ameliorated by the treatment of pexidartinib in vitro (**Fig. 2A-C**). In line with a phase 1/2 study of pexidartinib in relapsed/refractory FLT3-ITD+ AML,^18^ treatment of pexidartinb effectively suppressed the growth of MOLM-13 cells in vitro through reducing the phosphorylation of STAT5, STAT3 and AKT (**Fig. 2D-F**), reduced the leukemic burden and prolonged the survival of MOLM-13-xenografted mice (**Fig. 2G-I**).

**Figure 2.**
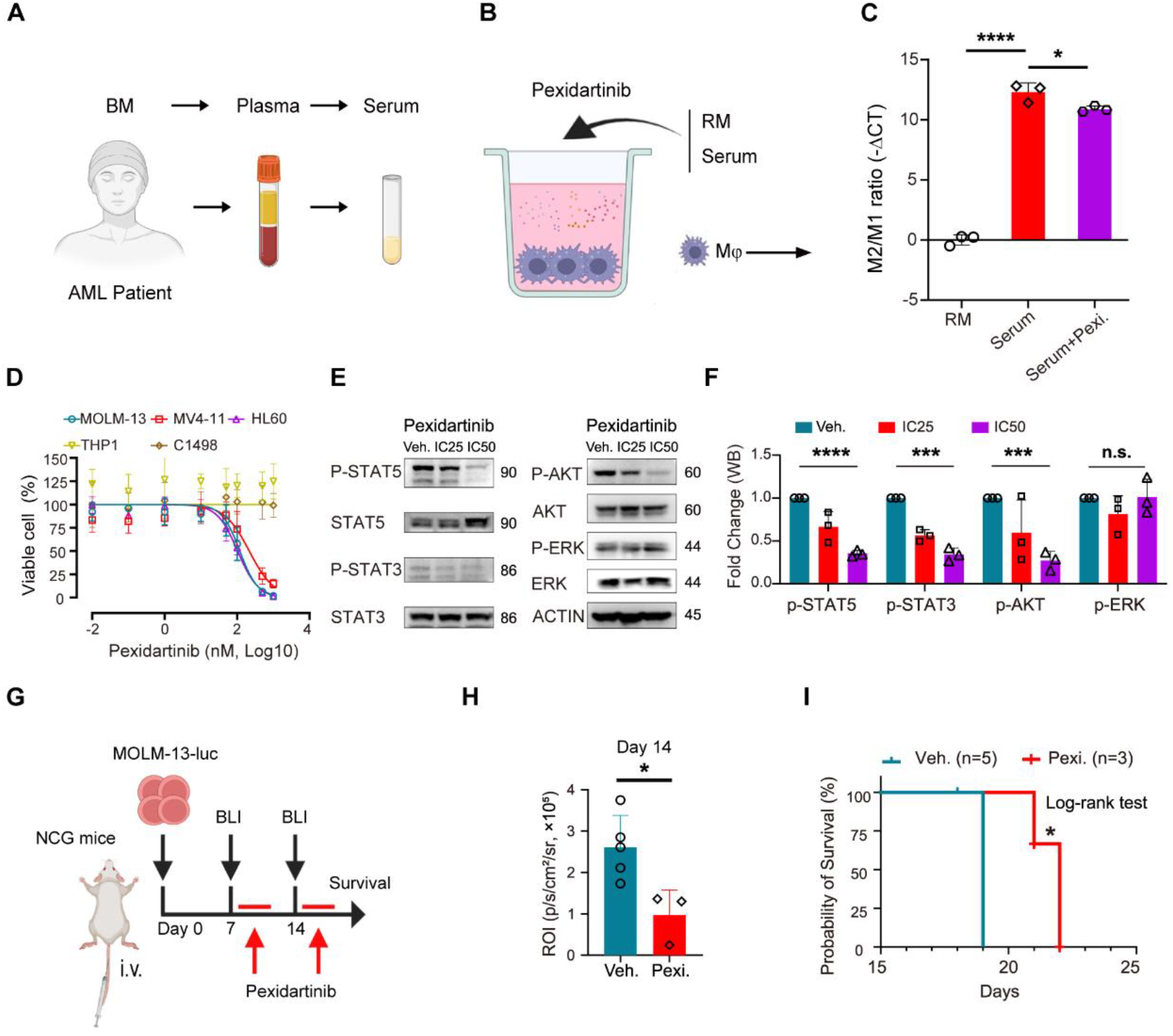
Targeting M2-LAM and FLT3-ITD+ AML by pexidartinib in vitro and in vivo. **(A-C)** Targeting M2-LAM in vitro. Serum from AML BM sample was derived (A) and added into THP-1-derived macrophages (B). RT-qPCR was performed and the polarization of macrophages was defined by the M2/M1 [-ΔCT(CD163-INOS)] ratio (C). **(D)** Dose-response curve of pexidartinib in MOLM-13, MV4-11, HL-60, THP-1and C1498 cell lines in vitro. **(E, F)** Western Blotting showing the expression and phosphorylation of FLT3 downstream signaling pathway in MOLM-13 cells after the treatment of pexidartinib (E). DMSO was used as vehicle control (Veh.). The band intensities were quantified by ImageJ and comparing after normalizing to the vehicle control (F). **(G-I)** Therapeutic efficacies pexidartinib in MOLM-13-xenografted mice. 6-8-weeks old immunocompromised NCG mice were injected with MOLM-13-luc cells (1×10^4^ per mouse) through tail vein. Bioluminescence imaging (BLI) was performed on day 7 and 14 after xenotransplantation. After confirmation of AML engraftment on day 7 by BLI, mice were treated with pexidartinib (oral gavage, 50 mg/kg) for two weeks (3 days per week) (G). Leukemic burdens (H) on day 14 were quantified based on BLI signals and compared (H) between vehicle (Veh.) and pexidartinb (Pexi.) group. The survival of animals (Veh. vs Pexi.) was recorded and compared based on log-rank test (I).

### mRNA-based FLT3L-CAR-Mφ showed enhanced phagocytic activities

It has been proposed that targeting FLT3 independent of its kinase activity may represent alternative therapeutic approaches for FLT3-ITD+ AML. In fact, FLT3 (also called CD135) is commonly overexpressed in the majority of AML cases (**Fig. 3A**) and represents an important immunotherapeutic target.^19^ FLT3-directed chimeric antigen receptor T (CAR-T)^13,14,20-22^ and CAR-NK^23^ immunotherapies are effective to target FLT3-ITD+ AML in preclinical animal models. Recent advances indicate that CAR-Macrophage (CAR-Mφ) exhibits enhanced and antigen-dependent phagocytosis of cancer cells in preclinical solid cancer models.^11,24-26^ However, the potential therapeutic efficacies of FLT3-directed CAR-Mφ in FLT3-ITD+ AML are still unknown.

**Figure 3.**
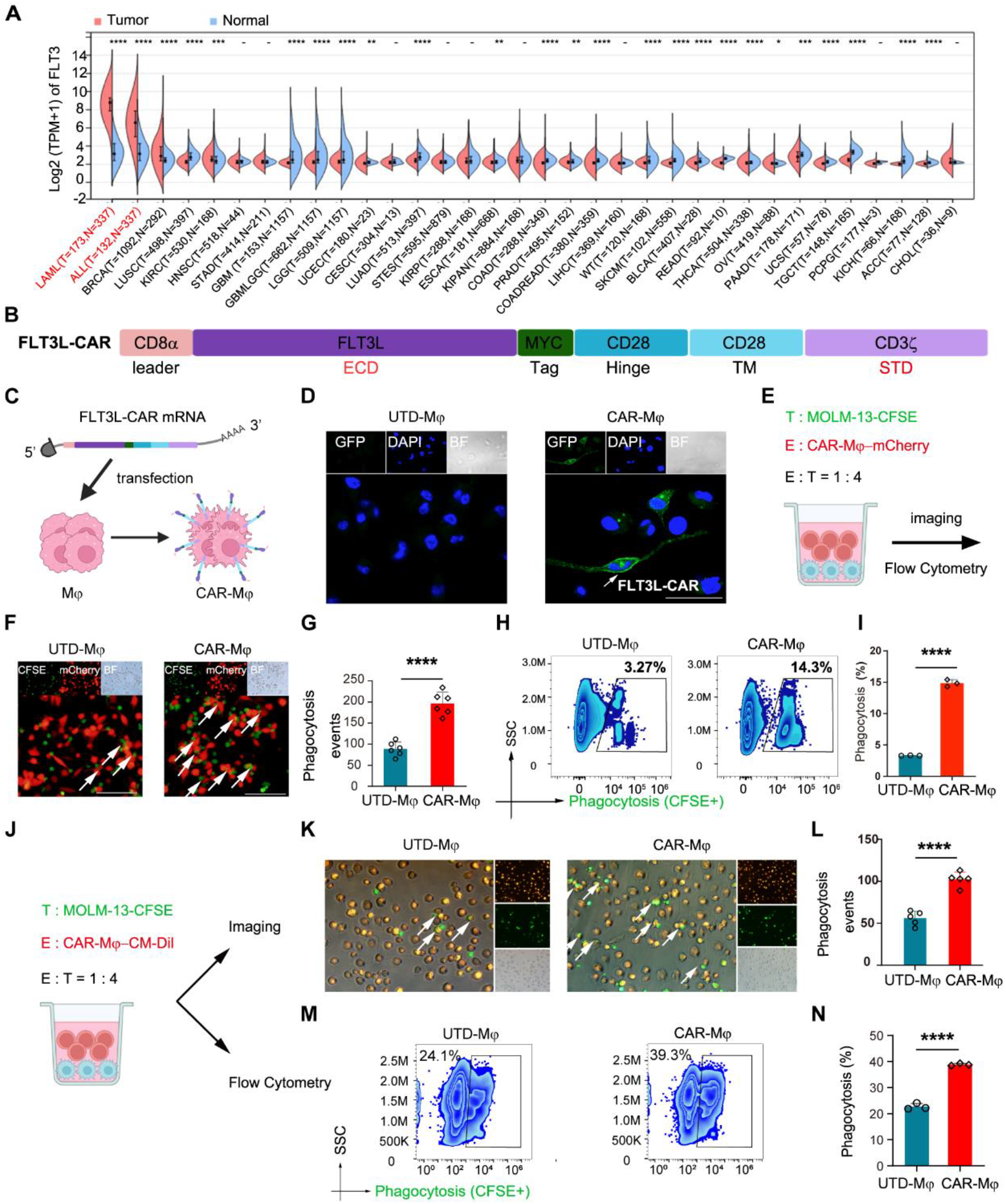
Design and characterization of mRNA-based FLT3L-CAR-Mφ. **(A)** Differential expression of FLT3 in human pan-cancers and their normal counterparts was analyzed by SangerBox (sangerbox.com) using TCGA and GETx database. **(B)** Design and components of FLT3-directed CAR-Mφ (FLT3L-CAR-Mφ). FLT3 ligand (FLT3L) and CD3ζ were used as extracellular domain (ECD) and signal transduction domain (STD) of CAR. CD28 hinge and CD28 transmembrane domain (TM) connected the FLT3L ECD and CD3ζ STD. A MYC-tag was inserted between the ECD and CD28 hinge. **(C-D)** FLT3L-CAR-encoding mRNA was in vitro transcribed and transfected in THP-1-deirved Mφ (C). The membrane expression of FLT3L-CAR was confirmed by anti-MYC immunofluorescence assay (D). **(E-I)** Phagocytic activities of mRNA-based FLT3L-CAR-Mφ-derived from mCherry+ THP-1 cells. FLT3L-CAR-Mφ (mCherry+) were cocultured with MOLM-13 cells (labelled with CFSE) for 48 hours and subjected for fluorescent imaging (F-G) and flow cytometry (H-I). Phagocytic events per field (F, arrows) were recorded and compared (G) in fluorescent imaging assay. The percentages of CFSE+ MOLM-13 cells in mCherry+ macrophages (H) were quantified in flow cytometry (I). Untransfected THP-1-derived macrophages (UTD) were used as control. **(J-N)** Phagocytic activities of mRNA-based FLT3L-CAR-Mφ-derived from umbilical cord blood mononuclear cells (UCBMNCs). UCBMNCs-derived macrophages were transfected with FLT3L-CAR-encoding mRNA, labelled with CM-Dil, and cocultured with CFSE-labelled MOLM-13 cells. The phagocytic activities of UCBMNCs-derived FLT3L-CAR-Mφ were measured by fluorescent imaging (K-L) and flow cytometry (M-N) and quantified as shown above. Untransfected UCBMNCs-derived macrophages (UTD) were used as control.

We then aimed to test whether FLT3-specific CAR-Mφ (FLT3L-CAR-Mφ hereafter) may show enhanced phagocytic activities to FLT3-ITD+ AML cells. FLT3 ligand and CD3ζ (**Fig. 3B**) were used as extracellular antigen-recognizing domain and intracellular signaling transduction domain of CAR as previously described, respectively.^11,14^ Transfection of THP-1 monocytic cells-derived macrophages with FLT3L-CAR mRNA (**Fig. 3C**) resulted in potent surface expression of FLT3L-CAR as demonstrated by immunofluorescent staining (**Fig. 3D, E**). MOLM-13 cells-expressing high level of FLT3^13,14^ were used as target cells for phagocytosis assays. Functionally, mRNA-based FLT3L-CAR-Mφ-derived from THP-1 cells showed enhanced phagocytic activities to MOLM-13 cells as demonstrated by increased proportion of double positive population by fluorescent imaging (**Fig. 3F, G**) and flow cytometry (**Fig. 3H, I**). Consistently, the enhanced phagocytic activities of mRNA-based FLT3L-CAR-Mφ were also observed in UCBMNCs-derived primary macrophages as shown by fluorescent imaging (**Fig. 3J-L**) and flow cytometry (**Fig. 3M, N**). These data indicate that FLT3L-CAR-Mφ are effective to target FLT3-ITD+ AML cells in vitro.

### Targeting FLT3-ITD+ AML by “off-the-shelf” FLT3L-CAR-Mφ

Generation of scalable and universal CAR-Mφ in a timely manner would be extremely crucial for the treatment of aggressive malignancies such as AML. CAR-expressing induced pluripotent stem cells (iPSC)^24^ and immortal THP-1 monocytic cells^11^ are two of the major sources to generate “off-the-shelf” CAR-Mφ. Comparing to iPSC,^24^ the differentiation of THP-1 cells into macrophages would be more flexible and time-effective (**Fig. 4A**). THP-1 cells-transduced with empty vector was used as control (EV-Mφ). After induction of macrophage differentiation by PMA for two days, surface expression of FLT3L-CAR was demonstrated by immunofluorescence assay (**Fig. 4B, C**). Comparing to EV-Mφ, “off-the-shelf” FLT3L-CAR-Mφ showed enhanced phagocytic activities to MOLM-13 cells as shown by an increased proportion of double-positive populations based on fluorescent imaging (**Fig. 4D-F**) and flow cytometry (**Fig. 4G, H**).

**Figure 4.**
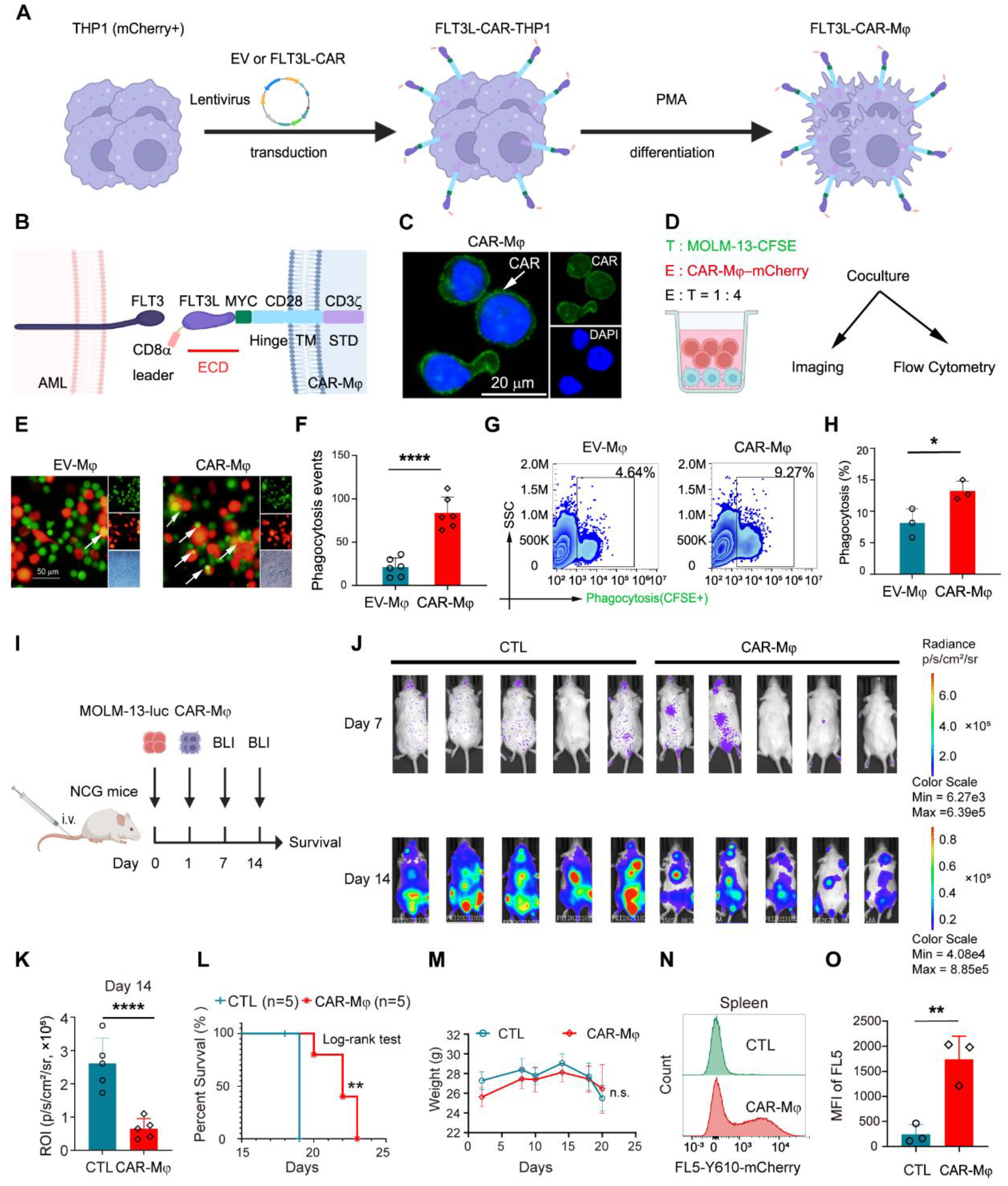
Efficacies of “off-the-shelf” FLT3L-CAR-Mφ in vitro and in vivo. **(A)** Scalable generation of “off-the-shelf” FLT3L-CAR-Mφ in vitro. Immortal THP-1 monocytic cells (mCherry+) were transduced with FLT3L-CAR-encoding lentivirus. FLT3L-CAR-expressing THP-1 cells were treated with PMA for two days to differentiate into universal and “off-the-shelf” FLT3L-CAR-Mφ. THP-1 cells-transduced with empty vector were also differentiated in macrophages and used as control (EV-Mφ). **(B, C)** Schematic diagram (B) and surface expression (C, anti-MYC immunofluorescence assay) of THP-1-derived “off-the-shelf” FLT3L-CAR-Mφ. **(D-H)** Phagocytic activities of “off-the-shelf” FLT3L-CAR-Mφ in vitro. FLT3L-CAR-Mφ (mCherry+) were cocultured with MOLM-13 cells (labelled with CFSE) for 48 hours (D) and subjected for fluorescent imaging (E-F) and flow cytometry (G-H). The phagocytic activities of “off-the-shelf” FLT3L-CAR-Mφ were defined by phagocytic events per field (E, white arrows) in fluorescent imaging (F) and percentages of CFSE+ cells within mCherry+ populations (G) in flow cytometry (H), respectively. **(I-O)** Therapeutic efficacies of THP-1-derived “off-the-shelf” FLT3L-CAR-Mφ in vivo. MOLM-13-luc cells (1×10^4^) were injected (day 0) into NCG mice via tail vein. MOLM-13-injecteed mice were treated with FLT3L-CAR-Mφ on day 1 and subjected for BLI on day 7 and day 14 (I). Mice-treated with PBS were used as control (CTL). BLI signal (J, K), survival (L) and body weights (M) of animals were recorded and compared. The infiltrations of mCherry+ FLT3L-CAR-Mφ into the spleen were detected by flow cytometry (N, O).

We then sought to test the therapeutic efficacies of FLT3-CAR-Mφ in AML xenograft mouse model. MOLM-13-luc cells (1 × 10^4^ per mouse) were transplanted into immunocompromised NOD/ShiltJGpt-*Prkd*^cem26Cd52^*Il2rg*^em26Cd22^/Gpt (NCG) mice via tail vein injection (**Fig. 4I**). Our pilot study revealed that injection of EV-Mφ did not show therapeutic effects comparing to saline control (CTL) (data not shown). Importantly, injection of FLT3L-CAR-Mφ (7 × 10^6^ per mouse) significantly reduced leukemic burden (**Fig. 4J, K**) and prolonged the survival of MOLM-13-xenografted mice (**Fig. 4L**). The body weight of FLT3-CAR-Mφ-treated animals are comparable to those from control groups (**Fig. 4M**), and no other side effects were observed. High percentage of FLT3L-CAR-Mφ (mCherry+) were detected in the spleen (**Fig. 4N, O**) of animal which MOLM-13 AML cells are highly infiltrated.

### Pexidartinib plus FLT3L-CAR-Mφ for the treatment of FLT3-ITD+ AML

FLT3-TKI plus induction chemotherapy represents standard-of-care for newly diagnosed FLT3-ITD+ AML ^27^. Interestingly, treatment of low dose (IC25 and IC50) FLT3-TKI (midostaurin, gilteritinib, crenolanib, sorafenib, quizartinib, and pexidartinib) resulted in an increased expression of glycosylated FLT3 (160 kDa) in FLT3-ITD+ MOLM-13 cells (**Fig. 5A**). The upregulation of FLT3 was also demonstrated in another FLT3-ITD+ MV4-11 AML cells (**Fig. 5B**). Of note, the effects of FLT3/CSF1R dual inhibitor pexidartinib^18,28^ were most significant. Indeed, pexidartinib-treated MOLM-13 cells showed increased surface localization of FLT3 as demonstrated by flow cytometry (**Fig. 5C, D**). Comparing to the vehicle control, exposure of pexidartinib sensitized MOLM-13 cells to the treatment of FLT3L-CAR-Mφ as shown by fluorescent imaging in vitro (**Fig. 5F, G**).

**Figure 5.**
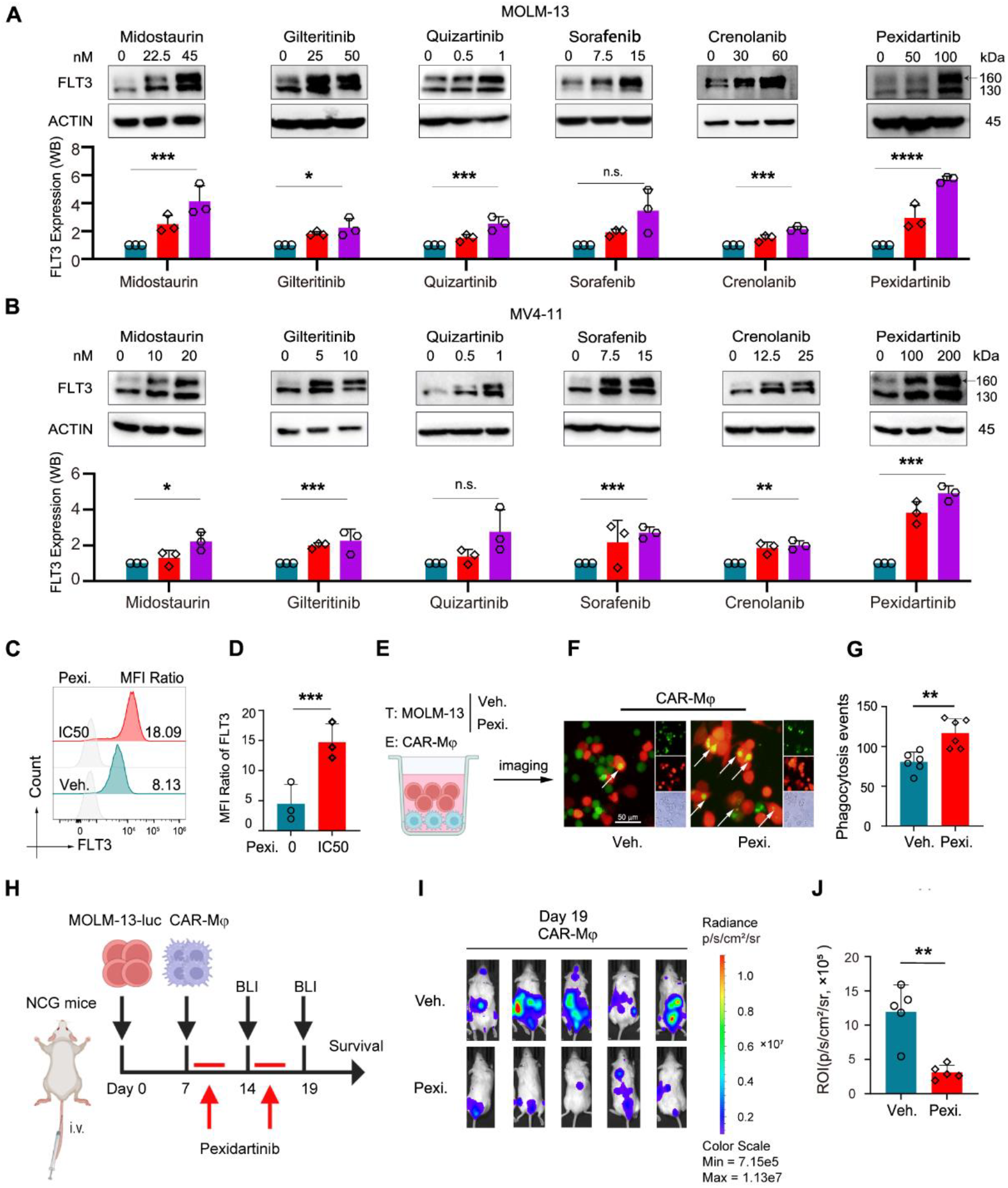
Pexidartinib sensitized FLT3-ITD+ AML cells to the treatment of FLT3L-CAR-Mφ in vitro and in vivo. **(A-B)** Western Blotting showing the expression of FLT3 upon treatment of FLT3-TKIs in MOLM-13 (A) and MV4-11 (B) cells in vitro. Relative FLT3 expression was quantified by normalizing to the corresponding ACTIN. **(C-G)** Exposure of pexidartinib upregulated surface expression of FLT3 (C-D, flow cytometry) in MOLM-13 cells in vitro and enhanced the phagocytic activities of FLT3L-CAR-Mφ in vitro as shown by coculture (E) and fluorescent imaging assays (F-G). (**H-J)** MOLM-13 cells (1×10^4^) were injected (day 0) into NCG mice via tail vein. MOLM-13-injecteed mice were treated with FLT3L-CAR-Mφ on day 7. FLT3L-CAR-Mφ-treated mice were then administrated with PBS control (Veh.) or pexidartinib (Pexi. 50mg/kg) for two weeks (3 days per week). BLI signals of animals on day 19 were recorded (I) and compared (J).

We then sought to test whether pexidartinib treatment may further enhance the therapeutic efficacies of FLT3L-CAR-Mφ in MOLM-13-xenografted mouse model. MOLM-13-luc cells (1 × 10^4^ per mouse) were transplanted into NCG mice via tail vein injection (**Fig. 5H**). 7-days-post transplantation, animals were first injected with FLT3L-CAR-Mφ (7 × 10^6^ cells per mouse, i.v.). Animals were divided into two groups randomly and treated with vehicle control or two cycle of pexidartinb (50 mg/kg daily, oral gavage, 3 days per week) (**Fig. 5H**). Comparing to vehicle group, administration of pexidartinib further reduced the leukemic burden of FLT3L-CAR-Mφ-treated animals (**Fig. 5I, J**), suggesting the synergistic effects of pexidartinib plus FLT3L-CAR-Mφ in vivo.

## Discussion

In this study, we provide novel insights into the roles of macrophages during the pathogenesis and therapeutics in FLT3-ITD+ AML. To the best of our knowledge, this is the first “proof-of-concept” study about the preclinical efficacies of FLT3-TKI plus CAR-Macrophage combination therapy for the treatment of FLT3-ITD+ AML in preclinical models.

First, our data indicate that FLT3/CSF1R dual inhibitor pexidartinib is effective to target M2-LAM and FLT3-ITD+ AML. Recent advances indicate that the immunosuppressive M2-LAM are highly infiltrated in human AML patients^8^ and preclinical AML animal models,^29^ drive leukemic transformation and progression,^29,30^ facilitate immune escape,^31^ and associated with inferior prognosis of AML patients.^8,32^ Consistently, here we show that conditioned medium from FLT3-ITD+ AML cells and serum from FLT3-ITD+ AML patients effectively polarize M2-LAM in vitro. Unexpectedly, M2-LAM effectively protected FLT3-ITD+ AML cells from the treatment of FLT3-TKI quizartinib through activating FLT3 signaling. In fact, both M2-LAM and AML cells express high level of CSF1R and inhibition of CSF1/CSF-1R reprograms M2-LAM^33^ and exhibit antitumor activity in AML.^34,35^ Our data reveal that FLT3/CSF1R dual inhibitor pexidartinb could suppress M2-LAM and target FLT3-ITD+ AML cells spontaneously. The therapeutic efficacies of pexidartinib in FLT3-ITD+ AML may be further tested in immunocompetent mouse model.

Second, out data reveal that FLT3-directed CAR-Macrophage is effective to target FLT3-ITD+ AML. Recent studies indicate that FLT3-ITD+ measurable residual disease (FLT3-ITD-MRD) detected by next-generation sequencing (NGS) in AML patients is associated with high-risk of relapse and shorter overall survival.^36-39^ CAR-Macrophage shows enhanced antigen-dependent phagocytic activities to cancer cells and exhibits excellent tissue distribution and penetration,^11,24,40^ making them a promising therapeutic approach to eradicate minimal residual diseases such as FLT3-ITD-MRD in AML. In this work, we demonstrate that single dose treatment of FLT3L-CAR-Mφ significantly reduced leukemic burden and prolonged the survival of AML xenografted immunocompromised mice without notable side effects in terms of body weight. The therapeutic efficacies of FLT3L-CAR-Mφ may be further explored in AML patient-derived xenograft (PDX) mouse model.

Third, our data suggest that pexidartinb plus FLT3-directed CAR-Macrophage combination therapy showed synergistic effects for the treatment of FLT3-ITD+ AML. Antigen loss and immune escape represent major obstacles for CAR-T- and CAR-NK-based cellular immunotherapy. Induction of antigen expression^22,41^ and epitope spreading (expansion of the immune response to secondary epitopes),^42^ design of dual antigen-targeted CAR,^43^ and combination of immune checkpoint inhibitors (ICI, such as CD47 antibody)^40^ are major strategies to overcome these obstacles. Here we FLT3/CSF1R dual inhibitor pexidartinib was most potent to induce the upregulation and surface localization of FLT3 in FLT3-ITD+ MOLM-13 and MV4-11 AML cells.

Importantly, pexidartinib sensitized MOLM-13 cells to the treatment of FLT3L-CAR-Mφ in vitro and in MOLM-13-xenografted mice. Dual antigen targeted CAR-Mφ may be further explored and the potential antigen spreading effects of FLT3L-CAR-Mφ may be investigated in peripheral blood mononuclear cells- or CD34^+^ hematopoietic stem cells-derived humanized mouse AML model.

In summary, our results provide novel insights into the pathogenic roles of M2-LAM during the leukemogenesis of FLT3-ITD+ AML. FLT3/CSF1R dual inhibitor pexidartinb suppresses M2-LAM, induces surface localization of FLT3 and sensitizes FLT3-ITD+ AML cells to the treatment of FLT3L-CAR-Mφ (**Graphical Abstract**). Pexidartinib plus FLT3L-CAR-Mφ could be a promising strategy for the treatment of FLT3-ITD+ AML.

## Author contributions

F.C. and B.L.H. conceived the study and designed the experiments. F.C., M.H.L., C.W.L., Y.Q.Z., Z.X.W., Z.Y., and J.Y.C. conducted the experiments. F.C., M.H.L., C.W.L., X.M.Y., J.B.X., L.W., and B.L.H. analyzed the data. F.C. and B.L.H. wrote, reviewed, edited, and revised the manuscript. B.L.H. directed and supervised the project. All authors discussed the results and approved the submission of the manuscript.

## Ethical approval and statement

All animal studies have been approved by the Ethical Committee at The Fifth Affiliated Hospital of Sun Yat-sen University and conformed to the ARRIVE guidelines. Informed consent was obtained from all subjects and the human studies were approved by the Institutional Review Boards from The Fifth Affiliated Hospital of Sun Yat-sen University. All experiments conformed to the principles set out in the WMA declaration of Helsinki.

## Data availability

The datasets used and/or analyzed during the current study are available from the corresponding author upon reasonable request.

## Acknowledgements

We are extremely grateful to all members (past and present) in HBL lab from Sun Yat-sen University. We thank Dr. Anskar Yu-Hung Leung, Cheuk-Him Man, Yiyue Zhang, Jun Chen, Xuan Sun, and Qing-Yun Wu for their generous support and insightful comments during the preparation and revision of manuscript. We appreciate for the technical support from Laboratory Animal Unit in The Fifth Affiliated Hospital of Sun Yat-sen University. This work was supported by grants from the National Natural Science Foundation of China (No. 32000569), the Basic and Applied Basic Research Foundation of Guangdong Province (No. 2019A1515110281), and Guangdong-Hong Kong-Macao University Joint Laboratory of Interventional Medicine (No. 2023LSYS001).

